# Tuning the mechanical properties of polymer-based surrogate materials for articular cartilage and vocal fold repair

**DOI:** 10.64898/2026.06.05.730092

**Authors:** Jessica Faber, Samuel Schlicht, Stefan Kniesburges, Anke Kaufmann, Lars Bräuer, Anna-Maria Liphardt, Maximilian Bachl, Tobias Pogarell, Matthias Stefan May, Michael Döllinger, Sarina K. Müller, Marcel Betsch, Mario Perl, Dietmar Drummer, Silvia Budday

**Affiliations:** Institute of Continuum Mechanics and Biomechanics, Department of Mechanical Engineering, Friedrich-Alexander-Universität Erlangen-Nürnberg, 90762 Fürth, Germany; Institute of Polymer Technology, Department of Mechanical Engineering, Friedrich-Alexander-Universität Erlangen-Nürnberg, 91058 Erlangen, Germany; Division of Phoniatrics and Pediatric Audiology, Department of Otorhinolaryngology, Head and Neck Surgery, University Hospital Erlangen, Medical School, Friedrich-Alexander-Universität Erlangen-Nürnberg, 91054 Erlangen, Germany; Institute of Functional and Clinical Anatomy, Friedrich-Alexander-Universität Erlangen-Nürnberg, 91054 Erlangen, Germany; Department of Internal Medicine 3 - Rheumatology & Immunology, University Hospital Erlangen, 91054 Erlangen, Germany; Institute of Radiology, University Hospital Erlangen, 91054 Erlangen, Germany; Imaging Science Institute, University Hospital Erlangen, 91054 Erlangen, Germany; Department of Otorhinolaryngology, Head and Neck Surgery, University Hospital Erlangen, Medical School, Friedrich-Alexander-Universität Erlangen-Nürnberg, 91054 Erlangen, Germany; Department of Traumatology and Orthopedics, University Hospital Erlangen, Friedrich-Alexander-Universität Erlangen-Nürnberg, 91056 Erlangen, Germany

**Keywords:** mechanical testing, articular cartilage, vocal folds, synthetic metamaterials, osteoarthritis, surrogate

## Abstract

The macroscopic biomechanical characteristics of soft and ultrasoft tissues, such as articular cartilage and vocal folds, significantly determine their physiological function. Treatments of widespread tissue degradations due to osteoarthritis in the knee or vocal fold impairment remain an unresolved challenge. For the design of implants for tissue repair after injury or disease, it is key to thoroughly understand the unique biomechanical properties of native tissues and potential substitute materials. We use multimodal mechanical testing methods combined with hyperelastic nonlinear continuum mechanics modeling, and finite element simulations to determine the macroscopic behavior of surrogate materials for human articular cartilage in the knee and human vocal folds. Our cyclic loading experiments reveal qualitative similarities for both tissues and their surrogates, including a nonlinear stress-strain behavior, hysteresis, and conditioning. We demonstrate the tunability of biomimetic and biosimilar stiffnesses of synthetic articular cartilage and vocal fold surrogates through tissue-specific process-material combinations. Our results demonstrate the feasibility of synthetic metamaterials in replicating essential passive biomechanical functions with great potential for future treatment options.

## 1 Introduction

### 1.1 Biomechanical function of soft tissues

Soft tissues, such as articular cartilage, as well as ultrasoft tissues, such as vocal folds, perform specific and in many cases vital biomechanical functions in the human body [Sophia Fox et al., 2009, Humphrey, 2003]. Degradation of these tissues can result from trauma, chronic mechanical stress, inflammatory diseases, degenerative processes, and cancer [Sophia Fox et al., 2009, Petitjean et al., 2023]. These conditions typically considerably impair their biomechanical function and the patients’ quality of life [Hunter and Bierma-Zeinstra, 2019]. Osteoarthritis, for instance, is a degenerative joint condition that is often initiated by cartilage damage and thus affects millions of individuals worldwide [GBD 2021 Osteoarthritis Collaborators, 2023, Long et al., 2022]. It is characterized by progressive cartilage degradation, subchondral bone remodeling, joint inflammation, and persistent pain [Hunter and Bierma-Zeinstra, 2019, Long et al., 2022]. Common symptoms alongside consistent pain include joint stiffness, reduced mobility, and functional limitations in everyday activities [Hunter and Bierma-Zeinstra, 2019, GBD 2021 Osteoarthritis Collaborators, 2023]. Complementarily, vocal fold impairments or even partial or complete loss due to injury, inflammation, paralysis, scarring, or tumor-related resection profoundly affect functions, such as speech, swallowing, and the protection of airways [Rosenthal et al., 2007, Jiang et al., 2000]. These conditions considerably disrupt the patients’ communication, social interactions, and overall well-being, and furthermore pose a continuous risk of suffocation [Jiang et al., 2000]. Currently available surgical treatments often result in only partial recovery of vocal function, persistent voice limitations, or the complete absence of phonation [Hansen and Thibeault, 2006]. In particular, many patients undergo multiple interventions with limited success [Hansen and Thibeault, 2006].

A key characteristic of soft tissues is found in their stress-strain relationships when exposed to external loads, which has been described as early as 1967 [Fung, 1967, Humphrey, 2003]. Unlike many engineering materials, soft tissues do not show a linear stress-strain response. Instead, they display a nonlinear strain-stiffening response [Fung, 1967, Schinagl et al., 1997]. This nonlinear behavior plays a decisive role in the adaptation to physiological loads and the overall tissue function under complex loading conditions. A prominent example is the characteristic damping behavior of articular cartilage [Humphrey, 2003]. In addition to the intrinsic nonlinearity, soft tissues typically show pronounced viscoelastic effects, i.e., hysteresis and recoverable conditioning during cyclic loading [Mow et al., 1980, Huang et al., 2001, Eschweiler et al., 2021]. This behavior is associated with the relaxation of stresses under constant loading [Mow et al., 1980]. Combined with anisotropic properties, it enables soft tissues to meet a wide range of biomechanical requirements, such as a high elastic resistance during the initial peak of loading followed by the slow relaxation of induced stresses on extended timescales [Wilson et al., 2005, Wang et al., 2003]. Articular cartilage shows a locally varying depth-dependent stiffness and viscoelasticity [Schinagl et al., 1997, Chen et al., 2001]. The pronounced rate-dependency of mechanical characteristics allows for the efficient distribution of physiological loads evenly across joint surfaces. Therefore, it is key to fulfill fundamental biomechanical functions in the human body [Mow et al., 1980, Lu and Mow, 2008]. The intrinsic viscoelasticity is significantly influenced by the histomorphology of hyaline cartilage, characterized by cell morphology, microstructural architecture, and the underlying molecular composition [Wilson et al., 2005, Korhonen et al., 2008]. In particular, McCreery et al. [McCreery et al., 2023] identified proteoglycans as the predominant drivers for the pronounced hyperelasticity in hyaline cartilage. Complementarily, vocal folds obtain their physiological function based on their ultrasoft tissue composition during their flow-induced periodic oscillations [Titze, 1988]. These consist of a layered structure formed by the epithelium, lamina propria, and underlying muscle [Titze, 1988, Chan and Titze, 1999]. Similar to articular cartilage, vocal folds show a macroscopically nonlinear and anisotropic behavior across a wide range of oscillation frequencies [Chan and Titze, 1999, Kelleher et al., 2013].

Current engineered tissues and available implant solutions exhibit certain limitations and, in particular, fail to reliably and adequately repair soft tissue defects [Gasik et al., 2018]. In the treatment of cartilage degradations, established methods, such as microfracturing and autologous chondrocyte implantation, typically result in the formation of fibrocartilage formation instead of hyaline cartilage [Mithoefer et al., 2009]. Existing approaches predominantly yield constructs with insufficient biomechanical properties, which may lead to persistent joint symptoms and an overall impaired joint functionality [Peterson et al., 2010, Gasik et al., 2018]. Notably, clinically used implants do not reproduce the viscoelastic properties and intrinsic material nonlinearity required for sustained physiological performance [Gasik et al., 2018].

In vocal fold reconstruction, synthetic implants that allow the replication of the physiological phonation do not yet exist [Ling et al., 2015, Hamilton and Birchall, 2017]. Early trials with bioengineered implants showed promising effects in tests with small animals, such as rabbits and minipigs, but have not yet been transferred to human patients [Shiba et al., 2016, Schlegel et al., 2023, Ling et al., 2015]. In particular, prevailing therapeutic options employ merely symptomatic approaches, such as electrolarynges for synthetic voice generation, but do not address the actual replication of vocal folds and their protective properties [Hamilton and Birchall, 2017].

Tissue engineered implants for different soft tissues face several challenges related to scaffold design, cellular organization, tissue maturation, differentiation, and the intrinsic anisotropy [Jelodari et al., 2022, Stampoultzis et al., 2021]. In particular, the prevalence of inhomogeneous cell distributions, an inadequate diffusion of nutrients, and the frequently limited mechanical reinforcement commonly yield implants that deteriorate under physiological loading conditions [Jelodari et al., 2022, Malda et al., 2004]. These limitations arise from competing requirements for scaffold structures [Moutos et al., 2007]. While scaffolds are required to optimize physiological conditions for chondrocyte proliferation and differentiation, physiological loads often exceed loading conditions that can be mechanically tolerated by such scaffolds, which in turn limits the adaptability of the resulting biomechanical characteristics [Moutos et al., 2007]. These inherent mechanical and biological shortcomings substantially limit the clinical effectiveness [Stampoultzis et al., 2021]. Therefore, there is an evident need for implant strategies that allow for the reproducible long-term stability of biomimetic implant mechanics [Stampoultzis et al., 2021, Fu et al., 2020]. Hence, while a variety of approaches for the biomimetic engineering of both cartilage and laryngeal tissue has been and is still investigated, limitations remain, especially regarding insufficient biomechanical characteristics [Fu et al., 2020, Hamilton and Birchall, 2017].

In an attempt to overcome these limitations, we address the engineering of artificial soft tissue implants that mechanically mimic native tissue. Through this approach, we intentionally do not focus on the replication of micro- and mesoscopic morphological tissue characteristics but instead concentrate on mimicking the macroscopic mechanical tissue response [Moutos et al., 2007, Fu et al., 2020]. This allows us to avoid morphology related limitations and to access a significantly increased range of biomechanical properties.

To span the artificial replication of macroscopic behavior of a broad stiffness range of tissues (Figure 1), we focus on the design of surrogate materials for ultrasoft vocal folds and comparatively stiff articular cartilage.

**Figure 1.**
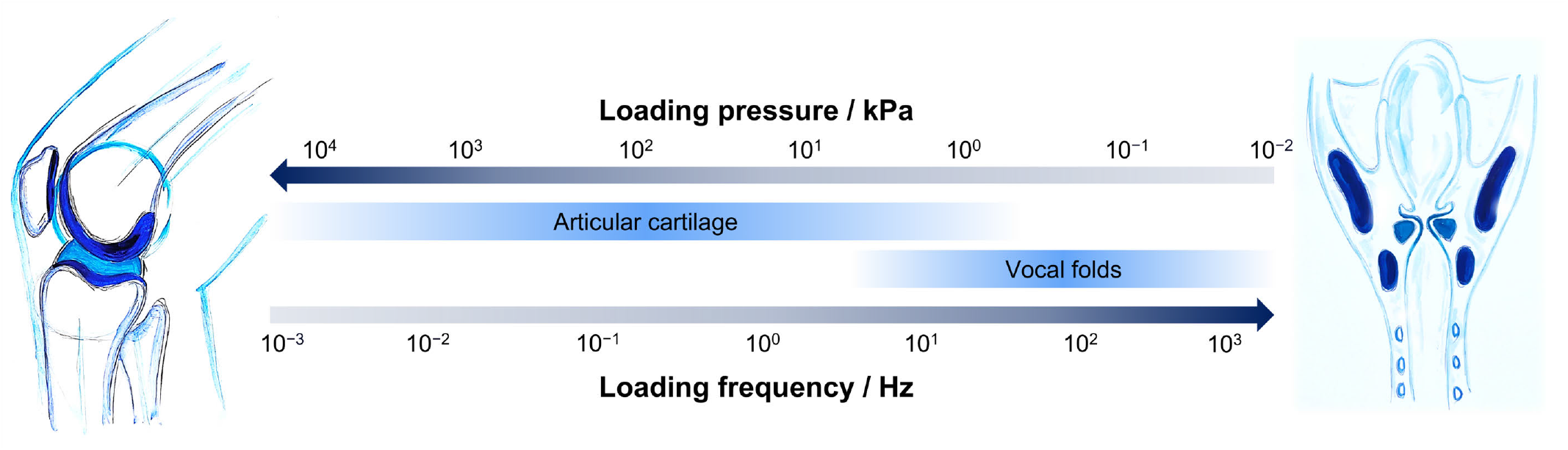
Loading conditions of different native tissues ranging from comparatively stiff human articular cartilage to ultrasoft vocal folds.

### 1.2 Approaches to biomimetic metamaterials

The preparation of biomimetic tissue surrogates necessitates the integration of (poro-)viscoelastic characteristics alongside their intrinsic nonlinearity. The synthetic replication of these complex, nonlinear biomechanics has been addressed in previous research based on a broad variety of process-material combinations. Based on non-reinforced hydrogels, early results by Metters et al. [Metters et al., 2000] showed the replication of effective moduli under compression exceeding 1 MPa while employing poly(ethylene glycol)-block-poly(lactic acid) (PEG-b-PLA) hydrogels with relative concentrations of 50% and 70%, respectively, while showing a pronounced degradation over time. Based on single-phase hydrogels, Nguyen et al. [Nguyen et al., 2012] showed the approximation of cartilage moduli through acrylate-cross-linked polyethylene glycol and demonstrated a significant variability of emerging moduli depending on the molecular mass of PEG precursors and their concentration. However, despite the temporary replication of elevated effective compressive moduli when applying increased hydrogel concentrations, several similar approaches on the engineering of cross-linked hydrogels reported merely reduced moduli [Bryant and Anseth, 2002, Roberts et al., 2011]. One frequently demonstrated approach that addresses the comparatively low stiffness of hydrogels is the additivation with fibrous or particulate reinforcements, which have been partially shown to achieve mechanical properties that approach those of native cartilage. Such an adaptability was exemplarily shown based on the addition of silk fibroin to metacrylated PEG hydrogels, yielding further increases in the effective moduli of prepared hydrogels [Fathi-Achachelouei et al., 2020]. In a cross-scale approach, Hu and Qu [Hu and Qu, 2019] showed the effect of micro- and nanoscale silica particles embedded in hydrogel-particle composites, with nanoscale particles yielding a significant stiffness increase, while no significant effects of particle size variations on a micro- and mesoscale could be identified. Alongside particle-modified systems, interpenetrating polymer networks (IPNs) have been shown to provide an alternative route to cartilage-like mechanics. A polyampholyte-based double network with a first network of poly(2-acrylamido-2-methylpropanesulfonic acid) and a second network of poly(N-isopropylacrylamide-co-acrylamide) obtained a compressive strength of approx. 25 MPa [Means et al., 2019], similar to cartilage tested under comparable strains. In this regard, the authors specifically stressed the qualitative comparability with cartilage biomechanics although significant quantitative limitations in mimicking the nonlinear stress-strain behavior under elevated strains were observed, which were particularly related to a relatively reduced nominal stress under compression.

Complementing the design of hydrogels for the replication of biomimetic mechanics, mesoscopically heterogeneous metamaterials have been applied for mimicking cartilage biomechanics. Chen et al. [Chen et al., 2025] applied mesoscopic, hyperboloid lattices for controlling emerging strains during tissue engineering of cartilage, albeit these showed a predominantly linear stress-strain behavior of applied scaffolds. Based on hybrid thermoplastic-hydrogel composites, nonlinear, cartilage-like stress-strain behaviors, despite showing comparably reduced effective moduli of swollen specimens, were furthermore demonstrated based on the variothermal powder bed fusion of polypropylene-agarose composites [Schlicht and Drummer, 2024c]. Further hybrid approaches for the generation of soft and ultrasoft metamaterials were described based on the laser-based consolidation and the subsequent biotechnological *in situ*-synthesis of bacterial nanocellulose, which were described to yield microporous composites with fiber diameters significantly below fiber dimensions usually obtained by technical fiber spinning processes such as melt electron writing [Schlicht et al., 2025b].

Complementing the generation of comparatively stiff cartilage-driven approaches, vocal fold models and surrogates have been predominantly described based on silicones and hydrogels. Among these, platinum-catalyzed, addition-curing silicones have been demonstrated to yield effective moduli in a range of 5 kPa - 90 kPa under quasi-static loading, which covers the mechanical range of both the epithelium and the embedded thyroarytenoid muscle. However, the specific replication of muscle tissue is intrinsically impeded by considerable variations in its stress-strain behavior in its relaxed and active state [Titze and Riede, 2010]. Despite such inherent limitations in the replication of variations in muscle biomechanics, homogeneous silicone models with intermediate stiffness in the range of about 10 - 30 kPa have been shown to self-oscillate in a range from 100 - 250 Hz under physiological flow. Elevated stiffnesses are associated with an increase in the glottal airflow resistance, hence increasing the subglottal pressure required for sustaining the oscillation, and furthermore increase the fundamental frequency [Luizard et al., 2023]. Beyond the passive function of vocal folds, the integration of ligament-like structures was shown to significantly increase obtainable frequencies, allowing to reproduce female vocal fold dynamics [Tur et al., 2023]. On a material level, Kazemirad et al. [Kazemirad et al., 2016] considered frequency-dependent characteristics of cross-linked hyaluronic acid - gelatin hydrogels, which corroborated a significant increase in measured elastic moduli at elevated frequencies, while the authors furthermore identified a pronounced reduction in the loss tangent at elevated, physiological frequencies, which is equivalent to a predominantly elastic behavior.

## 2 Materials and methods

### 2.1 Human articular cartilage specimens

Detailed information on the human articular cartilage specimens, including body donor characteristics, is provided in our previous study [Faber et al., 2026]. The obtained knee joints were frozen post-mortem and stored at *−* 20 °C. No significant differences in the mechanical properties between fresh and thawed articular cartilage after one freeze-thaw cycle are expected [Passman et al., 2011]. After slowly thawing of the joints at 8 °C for 2 to 3 days, Magnetic Resonance Imaging (MRI) measurements of a representative human knee joint (Donor no. 7, [Faber et al., 2026]) was performed using the Magnetom Vida 3-Tesla MRI scanner (Siemens Healthineers AG, Erlangen, Germany). Details on the MRI sequence data of the representative human knee joint can be found in Table A.1. Coronal slices were used for the anatomical visualization of the extraction position of a representative specimen from the lateral femoral condyle (Figure 2a).

**Figure 2.**
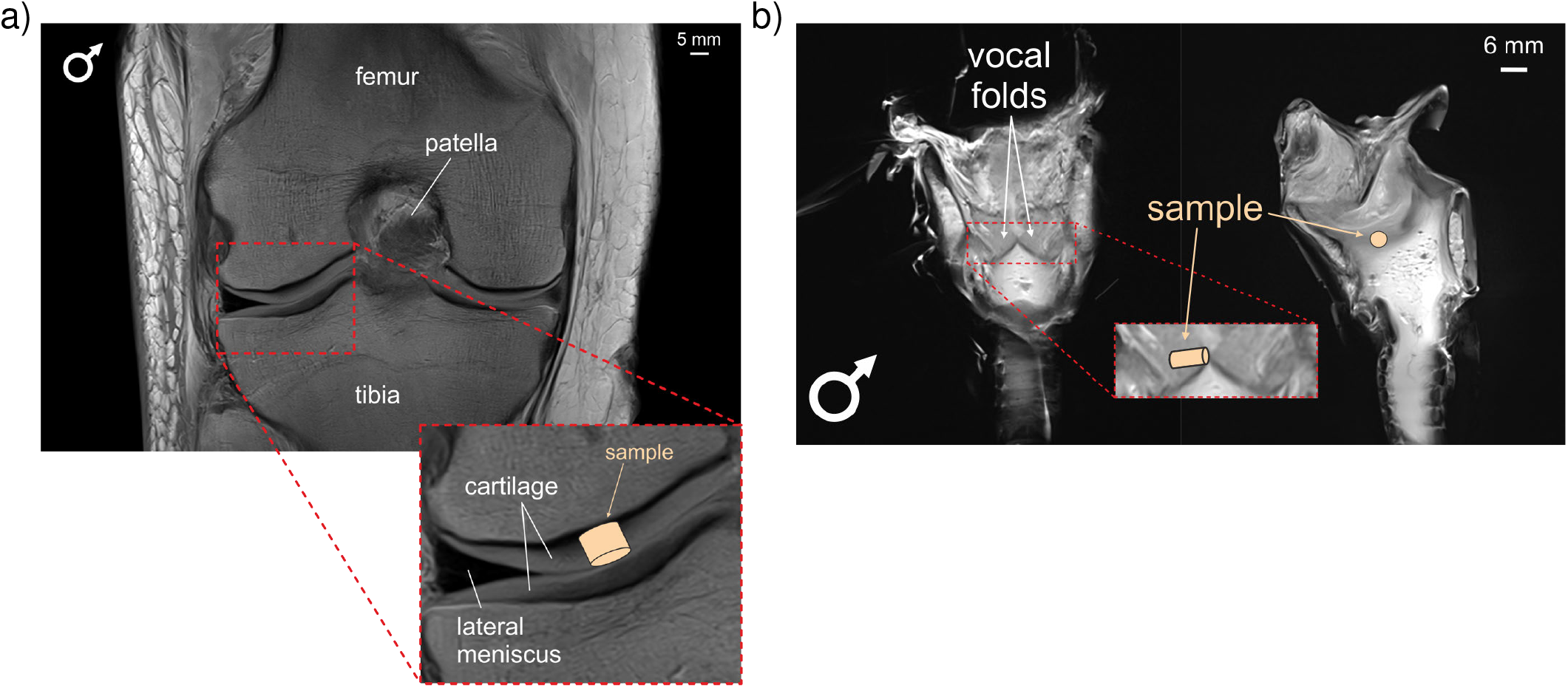
Schematic display of the position of a representative extracted cylindrical a) human articular cartilage, and b) human vocal fold specimens. Human articular cartilage samples from both sides (lateral and medial) of the femoral condyle and tibia plateau were extracted. One sample per vocal fold was extracted.

### 2.2 Human vocal fold specimens

Following the ethical approval (no. 405_18 B, Friedrich-Alexander-Universität Erlangen-Nürnberg, Germany), this study was conducted on the vocal folds of two larynges, one from a male and the other one from a female human body donor. The obtained larynges were frozen post-mortem and stored at − 20 °C. After thawing the larynges for approximately 3 h, MRI measurements of a representative human larynx (Donor no. 1, Table 1) was performed using the Magnetom Vida 3-Tesla MRI scanner (Siemens Healthineers, Erlangen, Germany).Details on the MRI sequence data of the representative human larynx can be found in Table A.2. Coronal and sagittal slices were used for the anatomical visualization of the extraction position of a representative vocal fold specimen (Figure 2b).

**Table 1.** Body donor information of larynges including the age, gender, and number of samples tested.

| Donor<br>no. | Age<br>in yrs | Gender | Number<br>of samples |
| --- | --- | --- | --- |
| 1 | 87 | male | 2 |
| 2 | 76 | female | 2 |
| total |  |  | 4 |

Afterwards, the vocal fold specimens were dissected and all mechanical tests were performed at 37 °C within 2 h. A summary of the body donor information of the larynges can be found in Table 1.

### 2.3 Human specimen preparation

The specimen preparation of articular cartilage from the knee joints are described in our previous study [Faber et al., 2026]. For specimen preparation of vocal folds, the larynges were cut into left and right halves until the vocal folds were accessible for specimen extraction and cylindrical specimens of the vocal folds and the underlying musculature were punched out using surgical punches with a diameter of 5 mm (Stiefel, Dublin, Ireland). All vocal fold specimens were tested immediately after sample extraction.

### 2.4 Surrogate specimen preparation

Surrogate materials were manufactured in cylindrical geometries with a diameter of 8 mm (articular cartilage from the knee joint) and 5 mm (vocal folds) in accordance with extracted tissue samples used for the biomechanical testing of native tissues. The generation of articular cartilage surrogates was based on the variothermal powder bed fusion [Schlicht et al., 2022, 2025a] of polypropylene–polyvinylpyrrolidone composites. A cryo-milled ethylene-propylene copolymer of type Ultrasint PP 1400 (Forward AM Technologies GmbH, Heidelberg, Germany) [Schlicht and Drummer, 2023, Mwania et al., 2025] and a polyvinylpyrrolidone (PVP) of type K90 (Carl Roth GmbH & Co. KG, Karlsruhe, Germany), exhibiting a nominal molar mass distribution from 900,000 g/mol to 1,200,000 g/mol, were used as received and blended in a laboratory mixer at 5000 /min for 10 min under the addition of liquid nitrogen for preventing overheating. Gravimetric fractions of 50 % and 55 % of PVP were applied. Based on the fractal, superposed exposure of prepared dry blends at ambient conditions (T = 25 °C), cylindrical specimens with a nominal height of 2 mm were manufactured, applying a combined volumetric energy density of 35 J mm^−3^ across five consecutive exposure steps, each consisting of four exposure sub-cycles. The applied process strategy followed a protocol described earlier [Schlicht and Drummer, 2024a,b] based on the fractal Peano curve, which enables the intermediate, discrete formation and crystallization of melt pools alongside the interlinked relaxation of residual stresses without inducing tensile stresses and warpage on the manufactured component. A constant laser power of 28 W, a hatch distance of 1.6 mm, and a layer height of 0.1 mm were applied. A counter-rotating roller was applied for powder coating to avoid the use of flow agents and to facilitate the processing of cryo-milled powders. To adapt locally occurring time-temperature profiles and the emerging structure formation, three distinct parameter variations were implemented based on varying the temporal distance of consecutive exposure cycles, yielding temporal distances of t = 2450 ms (Parameter set 1), t = 4900 ms (Parameter set 2), and t = 7350 ms (Parameter set 3).

Extracted laser-sintered specimens were subsequently stabilized in 20 wt.-% sodium persulfate solution (Carl Roth GmbH & Co. KG, Karlsruhe, Germany) through the thermal decomposition of persulfate ions and the interlinked formation of sulfate radicals, which, based on the intermediate formation of hydroxy radicals in aqueous media, induce the cross-linking of water-soluble PVP fractions [Gonzalez Ortiz et al., 2022]. The chosen concentration represents a stoichiometric excess to ensure the complete cross-linking of linear soluble PVP in accordance with Anderson et al. [Anderson et al., 1979]. Laser-sintered specimens were submerged at T = 23 °C for 300 s, followed by controlled heating (*dT/dt* = 2K*/*min) of the sodium persulfate solution to allow for simultaneous swelling and cross-linking of water-soluble PVP, yielding polyvinylpolypyrrolidone (PVPP). A PTFE-based lattice was used for fully submerging the laser-sintered specimens during the crosslinking to ensure complete coverage with the crosslinking agent. After reaching a temperature of 95 °C, specimens were kept at constant temperature for 4 h to ensure the complete decomposition of remaining persulfate. To ensure the removal of formed residual sulfate ions, obtained cross-linked specimens were washed and maintained in double-distilled water (T = 95 °C) for 12 h, followed by maceration in distilled water at room temperature for 96 h to yield an equilibrium state between water absorption and release (Figure 3, top).

**Figure 3.**
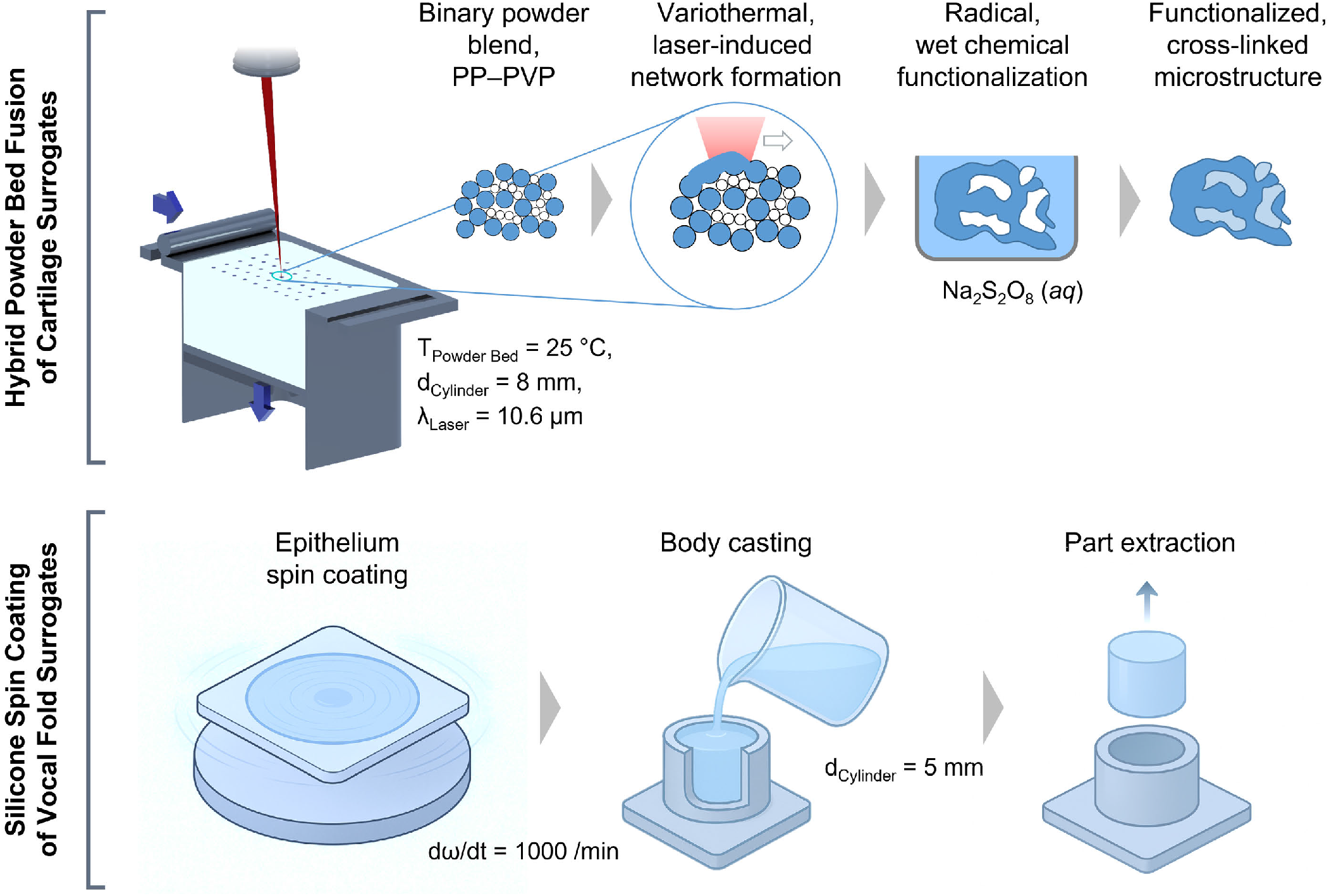
Schematic depiction of the fabrication process of surrogate materials based on hybrid powder bed fusion (cartilaginous tissue), and combined spin coating and casting (vocal fold tissue), fabrication not shown to scale.

The preparation of vocal fold surrogates was based on the addition-cross-linking silicone rubber of type SILPURAN 2420 (Wacker Chemie AG, Munich, Germany) that was modified with varying fractions of polydimethylsiloxane (PDMS) (Carl Roth GmbH & Co. KG, Karlsruhe, Germany) with a nominal viscosity of 5000 mPa s. Quantitative ratios of 1 : 1 : 8 and 1 : 1 : 7 (Curing component A : Curing component B : PDMS) were prepared through the casting of cylindrical specimens in additively manufactured separable cavities. To mimic the influence of artificial epithelium layers, addition-curing layers of non-modified silicone were prepared through the spin-coating of thin layers at a rotational speed of 1000 /min, yielding a thickness in the range of 20 *µ*m, following a previously described protocol [Kaufmann et al., 2024]. The applied protocol yielded cylindrical specimens with one-sided silicone layers in accordance with artificial larynx models with silicone-based epithelium analogs [Schlicht et al., 2024]. Obtained specimens were stored at T = 80 °C under inert conditions for 48 h prior to mechanical testing to ensure complete curing despite the dilution with inert PDMS. The complementary rotational molding of larynx models was based on geometries shown earlier by Tur et al. [Tur et al., 2023] (Figure 3, bottom).

Using axially symmetrical molds prepared by fused filament fabrication, artificial vocal fold surrogates were generated through rotational molding at varying rotational frequencies of 500 /min, 1000 /min, 1500 /min, and 2000 /min for obtaining defined variations in the thickness of emerging epithelium surrogates, following the protocol described by Kaufmann et al. [Kaufmann et al., 2024], with four rotational molding steps being conducted sequentially. Following the manufacturing of the silicone-based epithelium surrogate, a mixture of 1 : 1 : 7 (Curing component A : Curing component B : PDMS) was injected in the formed hollow part.

### 2.5 Experimental setup and data analysis

#### 2.5.1 Quasi-static mechanical experiments

For biomechanical testing, a Discovery HR-3 rheometer from TA Instruments (New Castle, Delaware, USA) mounted with an 8 mm parallel plate geometry was used to perform compression and tension experiments. To improve sample adhesion during testing, a circular piece of sandpaper with a diameter of 6 mm (K180, LUX Tools, Wermelskirchen, Germany) for human vocal folds specimens and corresponding surrogate material or with a diameter of 9 mm (K180, LUX Tools, Wermelskirchen, Germany) for most human articular cartilage specimens and corresponding surrogate material was glued to both geometries. Additionally, an immersion cup was attached to the lower plate. Next, the specimen was glued to the upper geometry using cyanoacrylate adhesive (Pattex superglue ultra gel, Henkel AG & Co. KGaA, Düsseldorf, Germany).

Subsequently, a thin layer of glue was applied to the lower plate and the specimen holder was lowered until contact of the specimen with the lower plate was visible. After about 60 seconds, the glue had cured and the previously attached immersion cup was filled with Dulbecco’s phosphate-buffered saline (DPBS) (Gibco Life Technologies, Thermo Fisher Scientific, Waltham, USA) until the entire specimen was covered in liquid. This procedure prevents structural degradation and dehydration during testing. All tests were performed at 37 °C to mimic *in vivo* conditions.

Initially, human articular cartilage samples were tested, as described in detail in our previous publication [Faber et al., 2026], and cyclic loading tests in compression up to 20 % strain and tension up to 2.5 % strain were performed with a loading rate of 40 *µ*m/s (average strain rate: 1.3 %/s), followed by stress relaxation experiments in both, compression and tension, with a loading rate of 100 *µ*m/s (average strain rate: 3.3 %/s) and a holding time of 300 s. Subsequently, the testing protocol was adapted and the loading rate for each test was adjusted to a constant strain rate of 1 %/s for cyclic loading and 2.5 %/s for stress relaxation tests. Experiments were performed using the adapted protocol for human vocal folds and SILPURAN-PDMS surrogates. Since PP-PVPP surrogates for human articular cartilage lacked sufficient stretchability up to % strain in tension, the protocol was modified and only tests in compression were performed.

#### 2.5.2 Dynamic mechanical experiments

Applying a sagittal elongation of 40 % ±2.1 relative to the length of the model vocal folds and volume flows of 25 and 30 standard liters per minute (slm), respectively, the emerging oscillation was characterized through high-speed imaging and the simultaneous recording of the sub-glottal pressure, following a protocol described earlier by Tur et al. [Tur et al., 2024]. A high-speed camera of type Phantom V2511 (Vision Research Inc., Wayne, NJ, USA) was used for recording, complemented by a subglottal pressure sensor of type XCS-93-5PSISG (Kulite Semiconductor Products, Inc., Leonia, NJ, USA). Subsequent analysis of obtained imaging data were based on the Glottis Analysis Tool [Kist et al., 2021].

### 2.6 Hyperelastic constitutive modeling calibrated with quasi-static experimental data

To characterize the elastic, time-independent behavior of native tissues and surrogate materials, the theory of nonlinear continuum mechanics is used, and the existence of a strain energy function Ψ is postulated. The strain energy function Ψ is decomposed into isochoric and volumetric parts

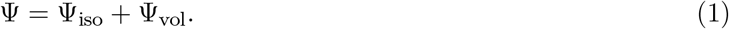

To capture the pronounced nonlinear behavior observed in native tissues [Budday et al., 2017, 2020, Faber et al., 2026], the modified one-term Ogden model [Ogden, 1972] for the isochoric response with its phenomenological, isotropic hyperelastic strain-energy function is formulated

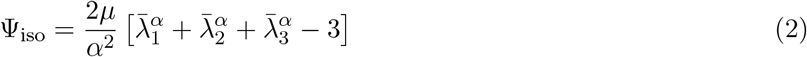

in terms of the nonlinearity parameter *α*, the classical shear modulus *µ*, and the isochoric principal stretches 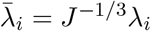 [Holzapfel, 2000]. For the volumetric response, Ogden [Ogden, 1972] proposed the generalized strain-energy function

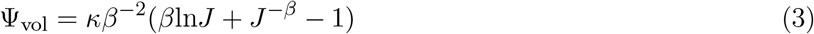

in terms of the bulk modulus *κ*, the empirical coefficient *β* and the Jacobian of the deformation gradient *J* = det(**F**). Based on the formulation of Simo and Miehe [Simo and Miehe, 1992], *β* is set to − 2, yielding the volumetric strain energy function

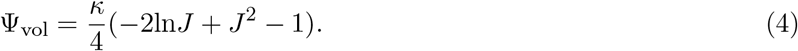

Since our experimental setup does not provide information on volumetric sample deformation during testing, the bulk modulus *κ* was computed from the classical shear modulus *µ* and the initial Poisson’s ratio *ν* using the relation

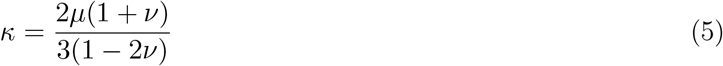

valid in the linear elastic regime. To circumvent numerical difficulties associated with incompressibility, near-incompressibility was enforced by setting *ν* = 0.49. Within the linear elastic regime, the (apparent) Young’s modulus can then be approximated to

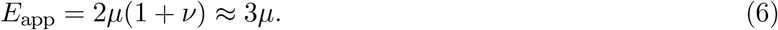

### 2.7 Data preprocessing

Experimental data from the unconditioned (first cycle) and conditioned (third cycle) responses of cyclic compression-tension tests on human tissues (articular cartilage and vocal folds) and SILPURAN-PDMS surrogates, as well as of the cyclic compression tests on PP-PVPP surrogates were used. To determine the hyperelastic material behavior, loading and unloading curves were averaged. Further details on the preprocessing procedure are provided in a previous study, which demonstrated that identifying parameters for each sample individually and subsequently averaging those parameters yields inaccurate results and can lead to nonlinear curves that deviate from the true averaged mechanical response [Hinrichsen et al., 2023].

### 2.8 Inverse parameter identification

Since each sample was glued to the upper and lower specimen holder, inhomogeneous deformation states during testing can be observed [Faber et al., 2022]. An inverse parameter identification scheme was applied to accurately capture the boundary conditions during testing and determine the classical shear modulus (*µ*) and the nonlinearity parameter (*α*) for the modified one-term Ogden model by coupling the finite element model with the trust reflective algorithm formulated by Branch et al. [Branch et al., 1999]. The optimal set of material parameters was identified for the first and third cycle of the averaged cyclic loading experimental data sets of human tissues and surrogate materials. Since the classical shear modulus (*µ*) is restricted to positive values, the optimization was constrained by (*µ >* 0). Further details regarding the inverse parameter identification scheme are provided in our previous study [Hinrichsen et al., 2023].

### 2.9 Microstructural investigations

Microstructural investigations of prepared surrogates were based on computed tomography, laser scanning microscopy, and scanning electron microscopy. Computed tomography was based on a sub-micron-CT, Fraunhofer Institute for Integrated Circuits (IIS) e.V., applying an isotropic spatial resolution of 5.0 µm. Complementary topographic investigations were based on a laser scanning microscope of type VK-X-1000 (Keyence Corp., Osaka, Japan). Microstructural investigations of silicone specimens relied on scanning electron microscopy (Zeiss Gemini, Zeiss Microscopy GmbH, Oberkochen, Germany) of cross-sections that were obtained through the symmetric cutting using scalpels and subsequent sputtering with Pd.

### 2.10 Statistical analysis

Statistical analysis of the maximum nominal stresses in compression (Figure 4a) and normalized nominal compressive stress (Figure 4b) of human tissue and surrogate materials was performed by one-way ANOVA followed by Tukey–Kramer tests for multiple comparisons if all samples were normally distributed and Kruskal-Wallis tests followed by Dunn-Sidak tests for multiple comparisons if not all samples were normally distributed using Statistics Toolbox in MATLAB version R2025b 25.2.0.3055257 (MathWorks, USA). A p-value lower than 0.05 was considered to be significant (*p < 0.05, **p < 0.01, ***p < 0.001, ****p < 0.0001).

**Figure 4.**
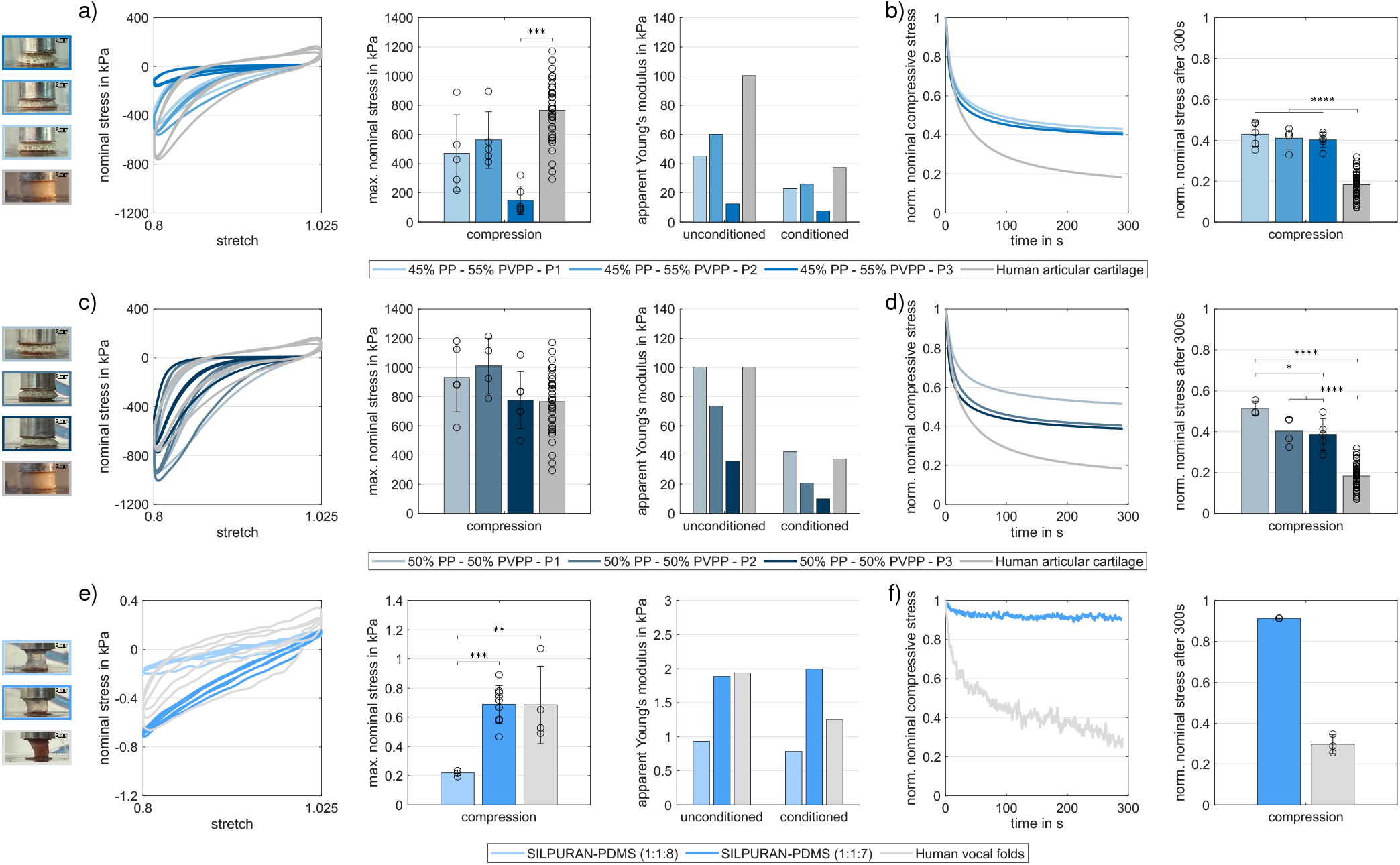
Mechanical characterization of human articular cartilage from knee joints and synthetic surrogates. Pictures show a representative glued sample of 45 % PP - 55 % PVPP (P1-P3), 50 % PP - 50 % PVPP (P1-P3), human articular cartilage, as well SILPURAN-PDMS composites with a quantitative ratio of 1:1:8, 1:1:7, and human vocal fold before testing. a) Cyclic loading behavior of 45 % PP - 55 % PVPP (P1: *n* = 5, P2: *n* = 5, P3: *n* = 6) and b) stress relaxation behavior of 45 % PP - 55 % PVPP (P1: *n* = 5, P2: *n* = 4, P3: *n* = 6). c) Cyclic loading behavior of 50 % PP - 50 % PVPP (P1: *n* = 5, P2: *n* = 4, P3: *n* = 6) and articular cartilage from human knee joints (*n* = 36, c.f. [Faber et al., 2026] Figure 2a-c) and d) stress relaxation behavior of 50 % PP - 50 % PVPP (P1: *n* = 3, P2: *n* = 4, P3: *n* = 5) and articular cartilage from human knee joints (*n* = 55, c.f. [Faber et al., 2026] Figure 2d, f). e) Cyclic loading behavior of SILPURAN-PDMS composites with quantitative ratios of 1:1:8 (*n* = 4), 1:1:7 (*n* = 9) and vocal folds (*n* = 4) and f) stress relaxation behavior of SILPURAN-PDMS composites with a quantitative ratio of 1:1:7 (*n* = 2) and vocal folds (*n* = 3). The stress relaxation behavior of all ultrasoft SILPURAN-PDMS composites with a quantitative ratio of 1:1:8 and 7 samples with a quantitative ratio of 1:1:8 are clearly unphysical (max. nominal compressive stress slightly above 1) and therefore not displayed as a result. This is strongly attributed to the low forces recorded during these tests, which lead to noise in the data. Due to the low number of samples (1:1:7 (*n* = 2) and vocal folds (*n* = 3)) for the quantification of the normalized stress relaxation behavior in compression, no statistical test for was applied. Significance values for Figure 4a-e: *p < 0.05, **p < 0.01, ***p < 0.001, ****p < 0.0001.

## 3 Results

### 3.1 Quasi-static mechanical behavior of human articular cartilage and PP-PVPP surrogates

Figure 4a and b show the macroscopic mechanical behavior under cyclic compression-tension and compression stress relaxation experiments of human articular cartilage and 45 % PP - 55 % PVPP surrogates with varying temporal distance of consecutive exposure cycles during the fabrication process. While incrementally reduced temporal delays between exposure cycles lead to a slight, insignificant decrease of the maximum nominal stress at a maximum strain of 20 % in compression from 561 kPa (Parameter 2) to 471 kPa (Parameter 1), extended delays (Parameter 3) further decrease it to 149 kPa. The 45 % PP - 55 % PVPP with reduced and moderate delays between exposure cycles (Parameter 1 and 2) show a similar mechanical response with slightly lower apparent Young’s moduli between 45 − 60 kPa for the unconditioned and between 23 *−* 26 kPa for the conditioned response compared to human articular cartilage with apparent Young’s moduli of 100 kPa for the unconditioned and of 37 kPa for the conditioned response (Table A.3, Figure A.1). Furthermore, 45 % PP – 55 % PVPP with reduced and moderate delays between exposure cycles (Parameter 1 and 2) show similar characteristics of articular cartilage including a nonlinear increase in stresses with increasing strains, substantial conditioning between the first and second loading cycle, and a pronounced hysteresis area, represented by the enclosed area between the loading and unloading curve. However, extended delays (Parameter 3) show a significantly softer response (***p=0.00012) with apparent Young’s moduli of 13 kPa for the unconditioned and of 8 kPa for the conditioned response (Table A.3, Figure A.1). Despite notable stress relaxation behavior up to around 60 % within 300 *s* of relaxation of 45 % PP – 55 % PVPP (independent of the temporal delays between exposure cycles), the surrogate still underestimates the stress relaxation of human articular cartilage by around 20 % points (****p<0.0001).

Based on these findings, we increased the concentration of PP and fabricated 50 % PP – 50 % PVPP surrogates to tailor the mechanical response toward the targeted mechanical behavior of human articular cartilage. Figure 4c and d show the macroscopic mechanical behavior under cyclic compression-tension and compression stress relaxation experiments of human articular cartilage and 50 % PP – 50 % PVPP surrogates with varying the temporal distance of consecutive exposure cycles during the fabrication process. In accordance with 45 % PP – 55 % PVPP surrogates, 50 % PP – 50 % PVPP surrogates depict a slight decrease of the maximum nominal stress at a maximum strain of 20 % in compression from 1010 kPa (Parameter 2) to 932 kPa for reduced temporal delays between exposure cycles (Parameter 1), and further decrease to 775 kPa for extended delays (Parameter 3). Independent of the temporal delays between exposure cycles, 50 % PP – 50 % PVPP surrogates are capable of mimicking the cyclic loading behavior of human articular cartilage, characterized by nonlinearity, conditioning and a pronounced hysteresis well. With reduced temporal delays between exposure cycles the stiffness can be tuned from 74 kPa (Parameter 2) to 100 kPa (Parameter 1) for the unconditioned and from 21 kPa (Parameter 2) to 42 kPa (Parameter 1) for the conditioned response (Table A.3, Figure A.1). But 50 % PP – 50 % PVPP surrogates still show less stress relaxation than human articular cartilage in compression, by around 20 % for moderate and extended temporal delays between exposure cycles (Parameters 2 and 3) and by around 30 % for reduced temporal delays between exposure cycles (Parameter 1) (****p<0.0001).

### 3.2 Quasi-static mechanical behavior of human vocal folds and SILPURAN-PDMS surrogates

Figure 4e and f show the macroscopic mechanical behavior of human vocal folds and SILPURAN-PDMS surrogates with quantitative ratios of 1:1:8 and 1:1:7. With a decrease in the fraction of PDMS from 8 to 7 parts with constant parts of the curing components A and B, the relative proportion between the curing components and PDMS increases. Consequently, SILPURAN-PDMS composites with a ratio of 1:1:7 have an increased cross-linking density and therefore show a significantly stiffer response during cyclic compression tests (***p=0.00055), achieving a maximum nominal stress of 0.7 kPa and a stiffness of 1.9 kPa, aligning with the maximum stress and apparent Young’s moduli for the unconditioned response of human vocal folds (Table A.3, Figure A.2). While human vocal folds show a pronounced nonlinear, hysteretic response during cyclic loading, with substantial conditioning between the first and the second loading cycle, SILPURAN-PDMS composites (1:1:7) exhibit a predominantly linear elastic behavior with negligible conditioning effects and hysteresis. As a result of the predominantly elastic behavior of SILPURAN-PDMS composites, the stiffness remains constant across all loading cycles. In contrast, the conditioned response of human vocal fold exhibits a lower stiffness, decreasing from 1.9 kPa to 1.3 kPa (Table A.3, Figure A.2). Furthermore, SILPURAN-PDMS composites with 1:1:7 relax up to 9 % within 300 *s* of relaxation and therefore show a less pronounced stress relaxation behavior than human vocal folds, which relax up to 70 % within 300 *s* of relaxation.

### 3.3 Dynamic mechanical behavior of human vocal folds and SILPURAN-PDMS surrogates

Figure 5 illustrates complementary oscillation experiments of PDMS-modified silicone surrogates under physiological volume flows. While quasi-static compression displays certain deviations between the native tissue and prepared surrogates with regard to their relaxation behavior, the intrinsically dynamic loading under oscillation indicates a less pronounced influence of relaxation phenomena on the oscillation and the interlinked phonation. The significant influence of the centripetal acceleration on emerging fundamental frequencies is in accordance with earlier observations, thus linking the emerging oscillation frequency to the vocal folds’ stiffness [Zhang et al., 2009]. Under reduced rotational frequencies, the centripetal acceleration applied during rotational molding shows a quasi-linear negative correlation with emerging fundamental frequencies and emerging subglottal pressures, respectively. Further increases in the applied rotational frequencies, however, yield a plateau, which is likely associated with the relatively increasing influence of the viscoelasticity of the embedded PDMS-modified body. While compression testing allows for the derivation of quantitative viscoelastic metrics of the ultrasoft body, the significant influence of the surrogate epithelium on emerging oscillation properties indicates the necessity to complementarily assess each surrogate under near-physiological, complex loading conditions.

**Figure 5.**
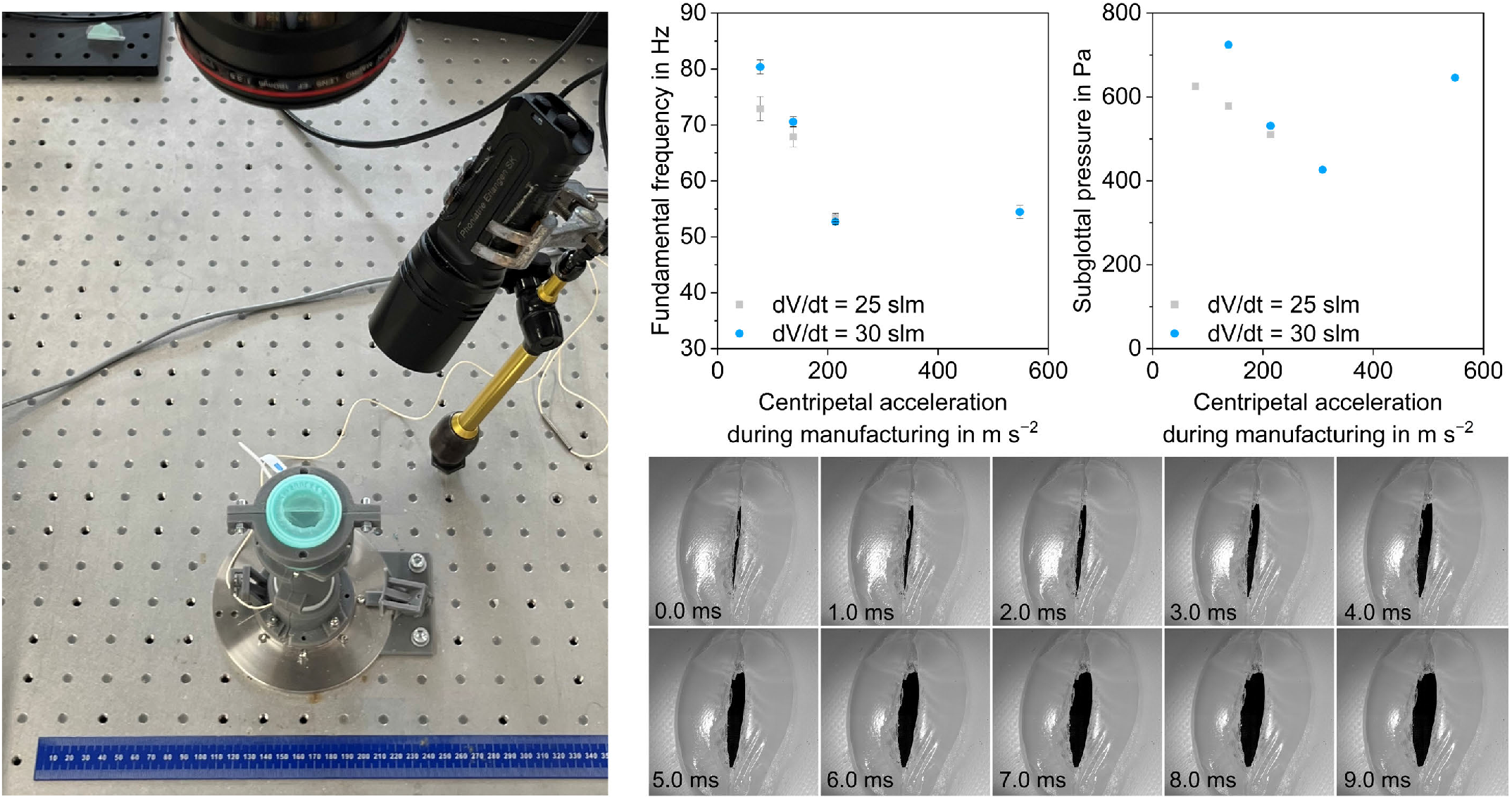
Oscillation characteristics of vocal fold surrogates with a quantitative ratio of 1:1:7 depending on manufacturing parameters under physiological volume flows, error bars display ±standard deviation based on n = 3.

Given the aforementioned influence of the epithelium, the quasi-static compression data should be interpreted in light of the load-bearing, anisotropic architecture of native vocal fold tissue. As a particular, anisotropic structure within the vocal folds, the vocal ligament is composed primarily of collagen and elastin fibers oriented along the anterior-posterior axis [Gray et al., 2000b,a]. These fibrous constituents exhibit a crimped, wavy configuration at rest and progressively straighten under increasing tensile strain, which is linked to the characteristic nonlinear stress-strain response observed in the lamina propria and in fiber-reinforced soft tissues in general [Gray et al., 2000b, Holzapfel et al., 2000]. As established in constitutive frameworks for fiber-reinforced soft tissues, collagen fibers are accordingly treated as tension-only elements that do not support compressive stresses, since the crimp structure impairs the mechanical load-bearing capability of buckling fibers under compression, which was described for arterial walls as a specific kind of fiber-reinforced soft tissue [Holzapfel et al., 2000]. Similar principles apply to the vocal ligament. Since the applied mediolateral compression acts orthogonally to the anterior-posterior fiber axis, the longitudinally oriented fibers can neither straighten nor carry load. The compressive response of the vocal fold is consequently governed by the soft, hydrated extracellular matrix of the lamina propria and the muscular body [Miri, 2014]. The homogeneous surrogate inherently approximates this compressive configuration, as it lacks oriented structures beyond the silicone-based epithelium. The observed agreement between surrogates and native tissue under quasi-static compression is therefore mechanically consistent: the vocal ligament does not contribute to the compressive response regardless of its anatomical presence.

The silicone-based surrogates successfully reproduced the self-oscillation and glottal flow aerodynamics consistent with established synthetic larynx models used for fluid-structure interaction analysis [Thomson et al., 2005]. In a recent review, Thomson [Thomson, 2024] classified the capabilities and limitations of synthetic, self-oscillating vocal fold models. Models were categorized into membranous and elastic solid architectures, the latter of which corresponds to the design philosophy of the surrogates presented herein. However, the fundamental frequencies (*f*_*o*_) recorded for our surrogates, which are found in the range of 50 to 80 Hz, are notably lower than typical male conversational speech at 100 to 120 Hz [Baken and Orlikoff, 2000]. This limitation is not unique to the present surrogates, but was shown earlier by Luizard et al. [Luizard et al., 2023], who reported a comparable frequency range of approximately 50 to 80 Hz for homogeneous silicone replicas fabricated from ultrasoft elastomers under varying pre-phonatory elongation, and identified the material stiffness as a primary determinant of *f*_*o*_ while observing only a marginal frequency response to longitudinal stretching in the absence of a fiber-reinforced ligament layer.

This limitation in the observed frequencies is mechanically consistent with the structural design of the current surrogates. Under phonation, longitudinal pre-strain imposed by the cricothyroid and thyroarytenoid muscles activates the vocal ligament in tension. The sequential step-by-step straightening of collagen fibers at increasing strain levels introduces a pronounced nonlinear stiffening mechanism, with the ligament exhibiting marked strain-stiffening above approximately 15 % tensile strain due to the previously discussed progressive collagen engagement [Chan et al., 2007, Gray et al., 2000a]. This mechanism is absent in our predominantly homogeneous surrogates. In human vocal folds, the vocal ligament accordingly serves as the primary tensile load-bearing component, which enables the exponential stress increase required to increase *f*_*o*_ during longitudinal tensioning [Titze and Hunter, 2004] towards physiological levels. The dynamic behavior of the oscillating body without ligamentous structures is hence influenced by the resulting stiffness and mass distribution. However, the isolated consideration of the vocal fold mass is insufficient [Titze, 2011] for determining the dynamic oscillation behavior. As demonstrated in material optimization studies for multi-layered vocal fold models, the precise tuning of layer geometry and stiffness is essential for achieving physiological displacement fields and frequency ranges [Schmidt et al., 2011]. In direct experimental support of the significance of the embedded ligament, Tur et al. [Tur et al., 2023] demonstrated that the integration of artificial ligament fibers of varying diameters and break resistances into multi-layered silicone vocal fold models enables a fundamental frequency range of approximately 200 to 520 Hz, which hence allows to cover both male and female phonation ranges. In particular, the authors showed in later work that the fiber tension represents the dominant parameter that governs the fundamental frequency, while the airflow rate primarily influences the subglottal pressure and oscillation amplitude [Tur et al., 2024, Gühring et al., 2024]. These effects correspond to the oscillation behavior in porcine full *ex vivo* larynx [Birk et al., 2017, Semmler et al., 2021] and *in vivo* hemilarynx canine experiments [Schlegel et al., 2024] where the oscillation frequency was found to be sensitive to the degree of adduction and elongation, which both increases the longitudinal tension within the fibrous ligament layer. Those influences were furthermore confirmed by Veltrup et al. [Veltrup et al., 2025] in an *ex vivo* human hemilarynx study. Hence, the surrogate employed in this study, which solely comprises a homogeneous silicone body alongside the silicone-based epithelium, lacks the ligament as a stiffening element. The behavior of non-reinforced surrogates thus effectively mimics a relaxed or passive state without the mechanical influence of the ligament under tensile loading, which indicates that, while homogeneous elastomers can replicate the onset of phonation, the inclusion of a distinct ligamentous layer is essential for accessing the higher frequency ranges that are associated with physiological phonation [Kniesburges et al., 2011, Thomson, 2024].

### 3.4 Microstructural characteristics of surrogates

Mechanical investigations of prepared surrogates indicate a significant influence of laser exposure parameters and the PP-to-PVPP ratio on the emerging rate-dependent properties of the composite surrogates. Complementarily, the cross-linking density of silicone surrogates significantly affects the stress-strain behavior. With regard to the interaction of mechanical and microstructural characteristics, representative surface and volumetric images exemplarily display the correlation of exposure histories and adapted PVPP fractions on the emerging internal architecture of manufactured specimens, displayed in Figure 6. Laser-optical micrographs obtained after superficial drying show a pronounced macroporosity alongside an exposure-dependent formation of the polypropylene network. In particular, the superficially observed network density decreases with an increasing delay between consecutive exposure cycles (Figure 6a), which is consistent with earlier investigations on the influence of time-temperature profiles on the formation of macroporous microstructures [Detsch et al., 2025]. Based on computed tomography, an overall reduction of the density of the formed polypropylene network at higher PVPP fractions (Figure 6b) could be observed. A comparatively reduced X-ray extinction in peripheral regions is consistent with partial drying of PVPP hydrogel during CT scans and the resulting formation of porous structures, whereas embedded PVPP hydrogel is preserved in central regions, which is morphologically represented as a more homogeneous core. These local variations suggest a gradient in the relative concentration of embedded hydrogels, which is furthermore in accordance with the wet chemical cross-linking sequence, in which an early dissolution of PVP near the surface may reduce the concentration of subsequently formed PVPP hydrogels in proximity to the surface of manufactured cylinders.

**Figure 6.**
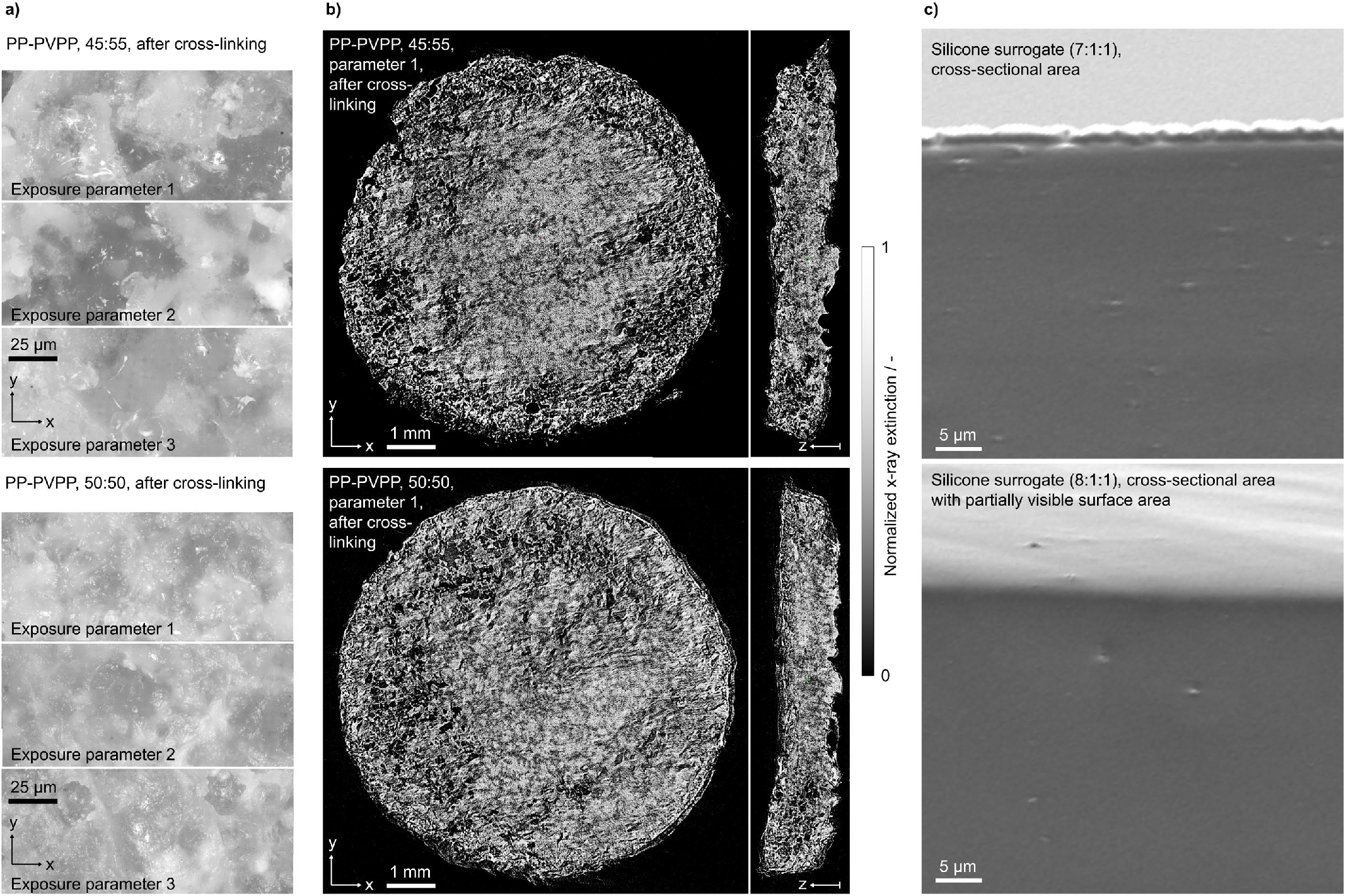
Microscopic depiction of exposure- and material-dependent topographic and morphological characteristics after cross-linking, showing a) the biphasic polyolefin-hydrogel surface structure; hydrogel fractions appear darker in reflected light micrographs, b) exemplary computed tomographic depiction of the stochastic composition of PP-PVPP composites after swelling, cross-linking and mechanical testing depending on the contained fraction of PVPP, and c) scanning electron micrographs of formed homogeneous cross-sectional microstructures of silicone-based vocal fold surrogates.

Elevated polyolefin network densities are correlated with increased normal stresses under compression. At equal PP and PVPP fractions (50-50), reduced temporal delays between exposure cycles (Parameter 1) promote a more efficient interparticle fusion and coalescence, inducing a stronger elastic contribution from the polypropylene network and a reduced viscous relaxation, whereas extended delays (Parameter 3) allow for tailoring the response toward the targeted biomimetic behavior, as observed in human articular cartilage. In this regard, computed tomography indicates the presence of merely incomplete load paths within the polypropylene network, displaying a stochastic network formation without a clear presence of oriented network structures perpendicular to the build plane in favor of a predominantly isotropic load-bearing PP network. An overview of scanning electron micrographs and corresponding g volumetric particle diameter distributions of PP and PVP can be found in Figure A.3.

In contrast to the intrinsically heterogeneous inner structure of laser-sintered hybrid composites, vocal fold surrogates exhibit a macro- and microscopically homogeneous morphology. In this regard, microscopic investigations corroborate the predominant absence of both macro- and microscopic pores, thus yielding a seamless transition between the spin-coated epithelium and the cast body (Figure 6c), which is consistent with the pronounced reproducibility of earlier quasi-static mechanical and function-related investigations.

## 4 Discussion

### 4.1 Qualitative mechanical similarities of human articular cartilage and human vocal folds

The complex mechanical characterization of native tissues under multiple loading modes and large strains is fundamental to thoroughly understand their mechanical properties [Faber et al., 2026] to design tissue-mimicking implants for tissue engineering applications and to develop tissue repair strategies [Weizel et al., 2020]. Tissue surrogates must satisfy specific mechanical criteria under physiological loading modes to be able to reproduce native function. Our study has demonstrated that the biomechanical characteristics of human articular cartilage and vocal folds exhibit considerable qualitative similarities, characterized by a nonlinear stress-strain response, in which stresses increase nonlinearly with increasing strains, substantial conditioning, where the initial loading cycle differs from the subsequent cycles, and a pronounced hysteresis with distinct loading and unloading paths whose enclosed area measures the dissipated energy during one loading cycle due to poroelastic and viscous effects. Furthermore, both tissues are biphasic materials, consisting of a layered, graded structure. Quantitatively, the maximum nominal stresses at 20 % compressive strain at comparable strain rates (1.3 %/s for human articular cartilage [Faber et al., 2026] and 1 %/s for human vocal folds) differ by a factor of around 10^3^, as well as the apparent Young’s moduli by a factor of around 50 for the unconditioned and by a factor of around 30 for the conditioned response (Table A.3, Figure A.1, Figure A.2). Despite substantial anatomic and stiffness differences, these shared intrinsic biomechanical characteristics are essential to fulfill physiological functions of both tissues. Moreover, they highlight the necessity of tissue-mimicking surrogates with similar qualitative but varying quantitative mechanical properties to replicate the macroscopic behavior of different tissues across diverse biomedical applications, from cartilage repair to vocal fold reproduction.

### 4.2 Morphological correlates of biomimetic mechanics

Through the consideration of morphologically divergent approaches to the replication of biomimetic soft tissue characteristics, we demonstrated the feasibility of synthetic pathways for the tunability of soft tissue surrogates. In this regard, we considered approaches that yielded both heterophasic and homophasic microstructures. Based on the variothermal powder bed fusion of polyolefin-hydrogel composites, we successfully demonstrated that a mesoscale polypropylene network, when interlinked with embedded hydrogels, exhibits a nonlinear, hysteretic stress-strain behavior including conditioning effects during cyclic loading as well as time-dependent stress relaxation behavior that likely rely on poro-viscoelastic effects. Complementarily, silicone-based replications of vocal fold mechanics provided indications that a homogeneous elastomer with controlled cross-linking density provides the general accessibility of ultrasoft biomechanics under the absence of meso- and microscopically heterophasic morphologies.

These findings allow us to identify the fraction of embedded hydrogels and, in particular, the density and intra-network connectivity of supporting, comparatively stiff polyolefin networks as predominant influences on the emergence of biomimetic stress relaxation. The replication of biomimetic stress-strain profiles, hysteresis and conditioning during cyclic loading is achievable through the control of emerging, stiffening networks, which we observed both on a mesoscale, as found for cartilage surrogates, and on a molecular scale, as observed for silicone-based vocal fold surrogates. In contrast, the replication of biomimetic relaxation properties requires the presence of embedded fractions that enable poro-viscoelastic effects, which may imply the predominant application of hydrogels. This interpretation has direct implications for future tunability. Within the cartilage-associated composite route, the temporal delay between consecutive exposure cycles alongside the fraction of embedded hydrogel precursor were found to jointly influence the mesoscale connectivity of fused polypropylene particles, which in turn influences the accessible volume of embedded hydrogel. Therefore, relaxation and hysteresis can be shifted toward an arbitrary biomechanical target behavior without loss of the nonlinear compression behavior. Within the addressed silicone-based route, the variation of the cross-linking density allows for the control of emerging stiffnesses. However, biomimetic relaxation behavior may necessitate the incorporation of fluid-containing domains to allow for the emergence of poro-viscoelastic mechanisms.

Complementarily, the morphologically represented variation in the local hydrogel density, observed through computed tomography, suggests that gradients in hydrogel content and network density may potentially be applied to tailor rate-dependent mechanical properties within a single part and to align these with locally varying mechanical requirements. Such gradients may provide a pathway to locally decouple the nonlinear stress-strain and stress relaxation behavior across different embedded materials. A further implication concerns the multi-objective performance across both compressive and tensile loading. In particular, prepared composites reproduce biomimetic compression characteristics of human cartilage but yield a significantly reduced effective modulus under tensile loading. These characteristics are consistent with the presence of incomplete tensile load paths in the stiff polypropylene network, which allow to transfer loads under compression but are associated with an insufficient stiffening under tensile loading. Morphology-related increases in the directed connectivity of the polyolefin network, for instance through the use of anisometric thermoplastic inclusions in the form of short fibers or platelets, may allow to significantly improve both the tensile stiffness as well as the surrogates’ performance under shear loading, while such structures could simultaneously preserve the fluid pathways that support stress relaxation in compression. In this way, stiffness placement, dissipation control, and tensile integrity under non-physiological conditions may be adaptable.

The two addressed approaches that strongly differ both in their microstructure and the underlying process strategies form a tunable spectrum, in which thermoplastics and ultrasoft elastomers represent its respective ends. The silicone-based approach obtains ultrasoft characteristics through a homogeneous elastic matrix, while the applied stochastic biphasic approach for the replication of cartilage biomechanics introduces time-dependent characteristics through embedded hydrogels and the implicit, process-based control of the network connectivity. The obtained findings thereby provide insights into the interactions of exposure parameters and the underlying material composition towards nonlinear rate-dependent mechanics. Hence, the control of the network density of both elastomers on a molecular level and the control of thermoplastic networks on a mesoscale level enables the biomechanics-driven tuning of metamaterials that allow us to replicate the passive mechanical behavior of soft tissues.

### 4.3 Limitations

A notable limitation of the present study is that all tissue samples were obtained from body donors and tested *in vitro*. Consequently, the specimens inherently reflect age-related changes, and the loading conditions applied during mechanical testing do not fully replicate the complex physiological environment experienced *in vivo*. In addition, the availability of human tissue is limited and therefore we could only test four vocal fold samples. Furthermore, the combined compression-tension testing of vocal fold specimens, applied during this study, employed strain rates below those of physiological phonation. This allowed us to maintain comparable conditions across tissues and surrogates, which supports the cross-tissue comparison and the comparison of corresponding surrogates, but limits the direct transfer to the highly dynamic *in vivo* behavior.

Concerning the generation of tissue surrogates, cartilage surrogates were based on statistical dry blends of cryo-milled particles with stochastic geometries. The transferability of obtained results may therefore be limited to materials of the used dimensions and geometries, since scale effects of thermoplastic network structures, formed after laser exposure, may alter emerging mechanics.

## 5 Conclusion

In the present study, we demonstrated the qualitative biomechanical similarity of cartilage and vocal fold tissue under combined compression-tension loading and the replication of these characteristics through synthetic metamaterials. Despite a significant difference in maximum stresses with a factor of around 10^3^ and in apparent Young’s moduli with a factor of around 50 for the unconditioned and with a factor of around 30 for the conditioned response, a conditioning-induced hysteretic behavior during cyclic loading and a pronounced relaxation of stress under compressive loading were observed for both cartilage and vocal folds. The engineering of metamaterials with corresponding biomimetic, tunable macroscopic mechanical characteristics was based on the variation of the density of thermoplastic networks and cross-linking densities of silicone networks, respectively. Relying on variothermal laser powder bed fusion, we could demonstrate the biomechanical tunability of polypropylene-hydrogel composites that exhibited a stochastic mesoscopic structure. Through the control of the polypropylene network density through adaptations in the fraction of hydrogel precursor, embedded during laser sintering, and the temporal variation of the laser exposure, we achieved apparent Young’s moduli between 13 and 100 kPa for the unconditioned and between 8 and 42 kPa for the conditioned response including biomimetic characteristics such as nonlinear, hysteretic stress-strain behavior with pronounced conditioning effects and a distinct normalized stress relaxation up to 60 %.

In addition, the replication of the ultrasoft biomechanics of the vocal folds was limited to initial compression with a stiffness of 1.9 kPa. While compressive properties could be replicated successfully, silicone-based surrogates were characterized by predominantly elastic properties in contrast to the pronounced viscous relaxation of native vocal folds, which underscores the significance of hydrogel-based materials and interlinked poro-viscoelastic properties for the replication of soft and ultrasoft tissue biomechanics. The accessibility of biomimetic mechanics through synthetic metamaterials thus motivates future investigations of material-process-immunology interactions as well as investigations into the ingrowth of subchondral tissue into open-porous metamaterials, such as the proposed PP-PVPP composites. Therefore, our investigations indicate the suitability of fully synthetic implants in biomechanically demanding, long-term treatments of degraded soft tissues.

## Author contributions

**Jessica Faber**: Methodology, Software, Validation, Formal analysis, Investigation, Data Curation, Writing - Original Draft, Visualization

**Samuel Schlicht**: Conceptualization, Methodology, Software, Validation, Formal Analysis, Investigation, Data Curation, Writing - Original Draft, Visualization

**Stefan Kniesburges**: Methodology, Investigation, Writing - Original Draft, Visualization

**Anke Kaufmann**: Investigation

**Lars Bräuer**: Methodology, Resources, Writing - Review and Editing **Anna-Maria Liphardt**: Investigation, Writing - Review and Editing **Maximilian Bachl**: Investigation, Writing - Review and Editing

**Tobias Pogarell**: Investigation, Writing - Review and Editing

**Matthias Stefan May**: Resources, Writing - Review and Editing

**Michael Döllinger**: Writing - Review and Editing, Resources, Funding acquisition

**Sarina Müller**: Writing - Review and Editing, Resources

**Marcel Betsch**: Methodology, Investigation, Writing - Review and Editing, Resources

**Mario Perl**: Methodology, Investigation, Writing - Review and Editing, Resources

**Dietmar Drummer**: Writing - Review and Editing, Supervision, Resources, Project administration, Funding acquisition

**Silvia Budday**: Conceptualization, Methodology, Resources, Writing - Original Draft, Supervision, Project administration, Funding acquisition

## Acknowledgements

The authors thank Jan Hinrichsen (Institute of Continuum Mechanics and Biomechanics, Friedrich-Alexander-Universität Erlangen-Nürnberg, Fürth, Germany) for implementing the initial inverse parameter identification code, Dr.-Ing. Bogac Tur (Dystonia and Speech Motor Control Lab, Department Otolaryngology - Head and Neck Surgery, Massachusetts Eye and Ear, Harvard Medical School) for his indispensable support in the experimental characterization of vocal fold surrogates, and Hannah Schlicht for her support in illustrating anatomical structures.

The authors wish to sincerely thank those who donated their bodies to science so that anatomical research could be performed. Results from such research can potentially improve patient care and increase mankind’s overall knowledge. Therefore, these donors and their families deserve our highest gratitude.

## Declarations

### Ethical approval

This study was approved by the Ethics Committee of Friedrich-Alexander-Universität Erlangen-Nürnberg, Germany, with the approval numbers 405_18 B and 18 − 405_2-B, and all procedures were conducted in accordance with the Declaration of Helsinki. Body donors had given their informed written consent to donate their body to research.

## Data Availability

Data will be made available on request.

## A Appendix

**Table A.1.** Sequence data from Magnetic Resonance Imaging (MRI) measurements of a representative knee joint (Donor no. 7, [Faber et al., 2026]).

| Sequence name | PD_TSE_fs_sag | PD_TSE_sag | PD_TSE_fs_cor | T1_TSE_cor |
| --- | --- | --- | --- | --- |
| Weighting | Proton Density | Proton Density | Proton Density | T1 |
| Imaging Technique | Turbo Spin Echo,<br>Fat Saturation | Turbo Spin Echo | Turbo Spin Echo,<br>Fat Saturation | Turbo Spin Echo |
| Plane of Acquisition | sagittal | sagittal | coronal | coronal |
| Repetition time (TR; ms) | 3400 | 2880 | 3040 | 666 |
| Echo time (TE; ms) | 40 | 33 | 31 | 10 |
| Voxel size (mm<sup>3</sup>) | 0.1953 x 0.1953 x 3 | 0.1736 x 0.1736 x 3 | 0.2131 x 0.2131 x 3 | 0.1465 x 0.1465 x 3 |
| Field of view (FOV; mm) | 149 x 149 | 149 x 149 | 150 x 150 | 150 x 150 |
| Number of Columns/Rows | 768/768 | 864/864 | 704/704 | 1024/1024 |

**Table A.2.**
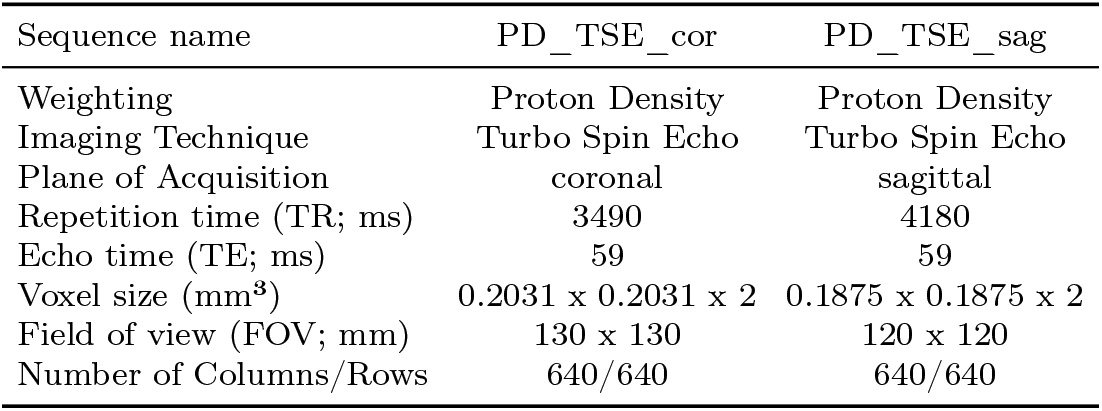
Sequence data from Magnetic Resonance Imaging (MRI) measurements of a representative larynx (Donor no. 1, Table 1).

**Table A.3.**
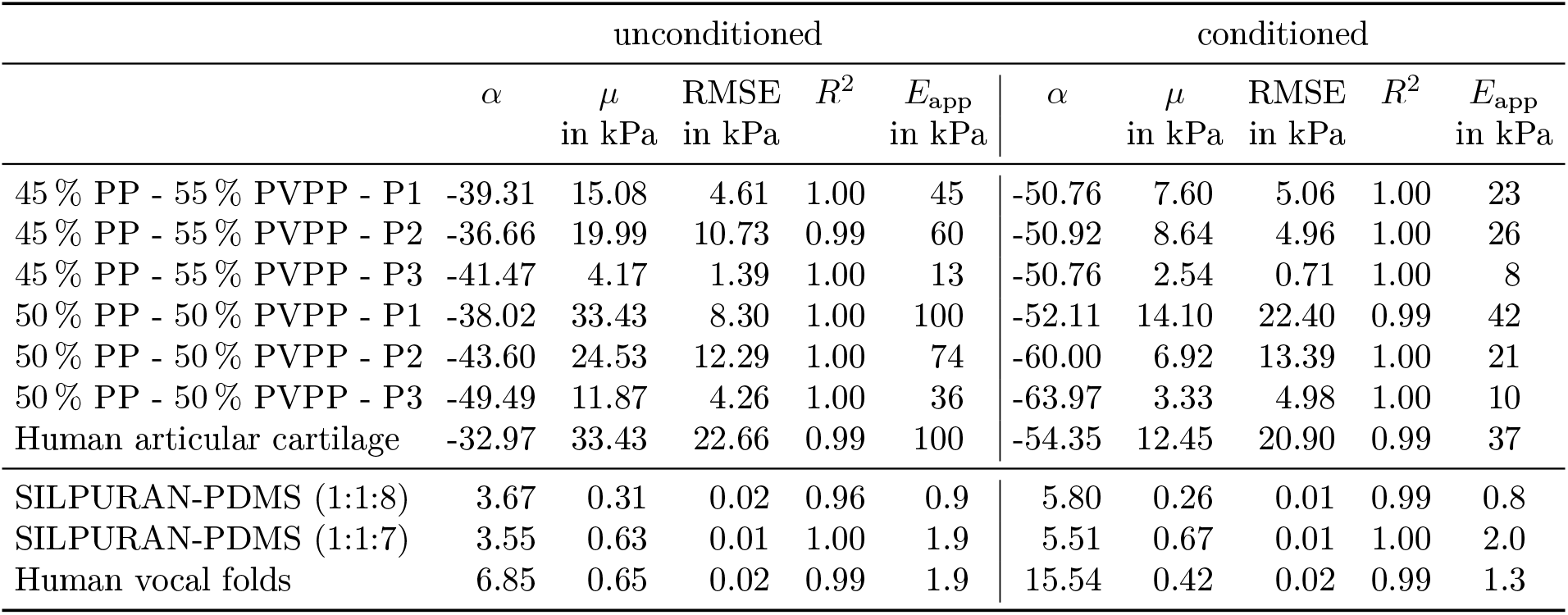
Nonlinearity parameters (*α*), classical shear moduli (*µ*), root mean square errors (RMSE), coefficients of determination (*R*^2^) and apparent Young’s moduli (*E*_app_) for the unconditioned and conditioned mechanical responses of 45 % PP - 55 % PVPP (P1: *n* = 5, P2: *n* = 5, P3: *n* = 6), b) 50 % PP - 50 % PVPP (P1: *n* = 5, P2: *n* = 4, P3: *n* = 6) and human articular cartilage (*n* = 36) obtained from fitting the modified one-term Ogden model to the first and third cycles of cyclic compression up to a maximum strain of 20 % in compression at a strain rate of 0.01/s (PP-PVPP) and 0.013/s (human articular cartilage) and of SILPURAN-PDMS composites with quantitative ratios of 1:1:8 (*n* = 4) and 1:1:7 (*n* = 9) and human vocal folds (*n* = 4) obtained from fitting the modified one-term Ogden model to the first and third cycle of cyclic compression-tension up to a maximum strain of 20 % in compression and 2.5 % in tension at a strain rate of 0.01/s.

**Figure A.1.**
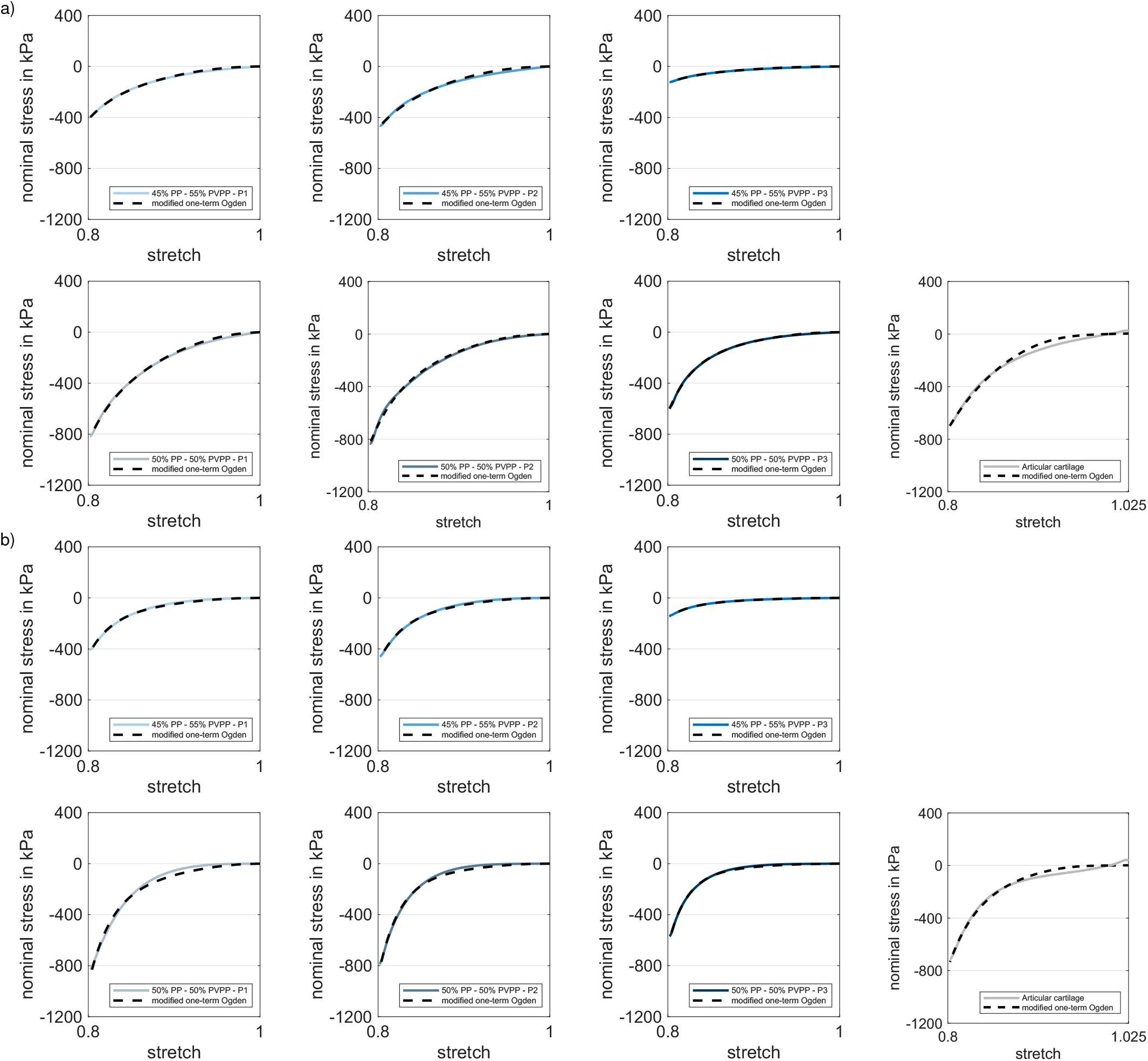
: Modified one-term Ogden model calibrated with the average a) unconditioned and b) conditioned mechanical responses of 45 % PP - 55 % PVPP (P1: *n* = 5, P2: *n* = 5, P3: *n* = 6), b) 50 % PP - 50 % PVPP (P1: *n* = 5, P2: *n* = 4, P3: *n* = 6), and articular cartilage from human knee joints (*n* = 36) during cyclic compression up to a strain of 20 % at a strain rate of 0.01/s (PP-PVPP) and 0.013/s (articular cartilage).

**Figure A.2.**
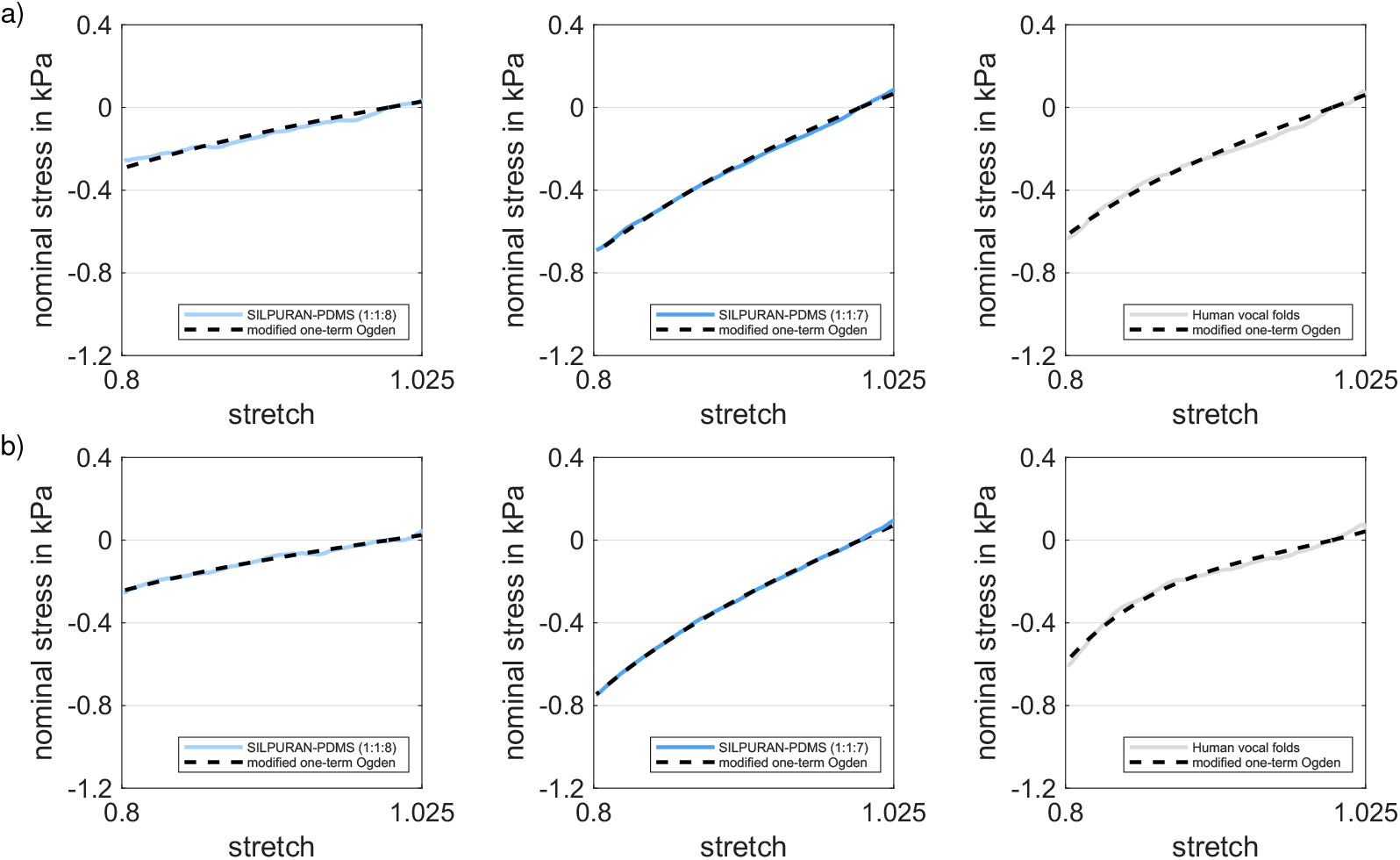
Modified one-term Ogden model calibrated with the average a) unconditioned and b) conditioned mechanical responses of SILPURAN-PDMS composites with quantitative ratios of 1:1:8 (*n* = 4), 1:1:7 (*n* = 9), and human vocal folds (*n* = 4) during cyclic compression-tension up to a maximum strain of 20 % in compression and 2.5 % in tension at a strain rate of 0.01/s.

**Figure A.3.**
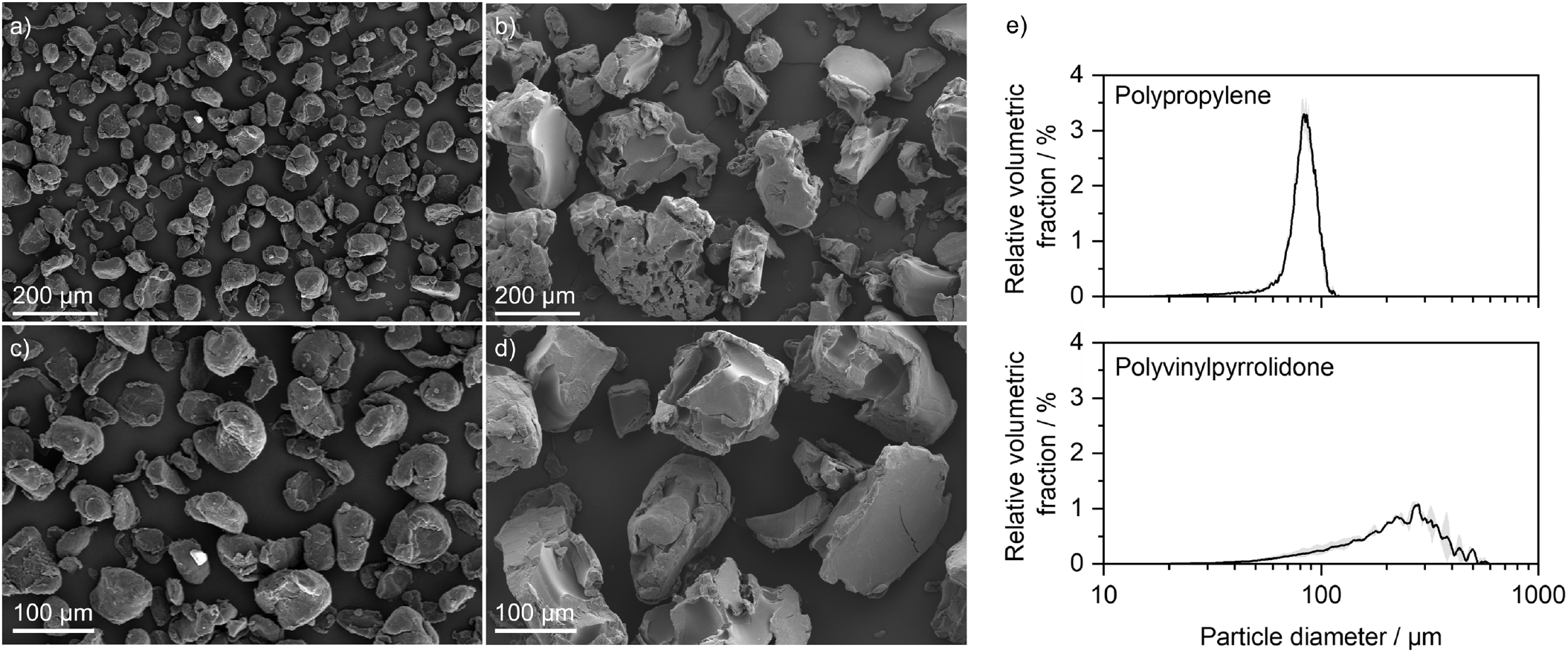
Overview of scanning electron micrographs of a), c) polypropylene (PP) and b), d) polyvinylpyrrolidone (PVP), used for the laser-based processing of articular cartilage surrogates, and e) corresponding volumetric particle diameter distributions.

